# Food-web architecture governs when predator advantage supports collective persistence

**DOI:** 10.64898/2026.05.19.726304

**Authors:** Ankit Vikrant, Serguei Saavedra

## Abstract

A recurring question across biological systems is when gains accrued by one part of a system also benefit the whole, and when they instead impose a collective cost. In ecological communities, consumers can increase their energetic gains through trophic interactions, yet those same interactions also determine whether all species persist. Here we show that food-web architecture governs whether predator advantage supports collective persistence, and that omnivory is a key condition under which the two diverge. Using a Lotka– Volterra-type food-web model formulated in terms of energy fluxes, we compare predator output power with the probability of feasibility, which quantifies the range of growth conditions compatible with positive coexistence. In two-species systems, these objectives show no generic alignment. In trophic chains, by contrast, increasing encounter rates makes predator advantage and coexistence mutually reinforcing. Basal omnivory reverses this pattern by shifting the power optimum towards the boundary of coexistence, where the intermediate consumer is lost. This pattern persists in larger networks, under heterogeneous encounter rates, and with saturating functional responses. Our results identify food-web architecture as the determinant of whether local energetic advantage scales up as collective persistence or instead becomes a coexistence cost.

## Introduction

A recurring question across biological systems is whether optimization at one level of organization benefits the system as a whole or instead imposes a collective cost. In ecological communities, this tension is especially acute. Consumers can increase their energetic gains through stronger or more numerous trophic interactions, yet those same interactions also determine whether all species can persist (May (1972); McCann et al. (1998); McCann (2000); Thébault & Fontaine (2010)). Understanding when local energetic advantage aligns with collective persistence is therefore central to explaining how food-web structure shapes ecological outcomes (McCann (2000); Thébault & Fontaine (2010)).

A classical framework for this problem is the maximum power principle, developed around the same time as the predator–prey Lotka–Volterra model (Lotka (1925)). The principle states that, under competition, configurations that capture and transform usable energy at higher rates are more likely to prevail. Lotka first framed this idea in terms of organismal fitness, arguing that fitness is enhanced by greater energetic output toward growth, reproduction, and maintenance (Lotka (1922a)). Later work generalized it into a broader principle of self-organization, potentially applicable across levels of biological organization and even beyond biology (Lotka (1922b); Odum & Pinkerton (1955)). Yet whether local energetic optimization promotes or erodes persistence at the scale of the ecological community remains unresolved (Lenton et al. (2021)).

In open systems, output power can be defined as the energy flux through a system component when energy enters through an external input channel. In ecosystems, solar energy entering through autotrophs provides such an input and is subsequently transformed into energetic fluxes across higher trophic levels. Maximizing output power for one or a few species may therefore select particular ecosystem states, each associated with a different set of surviving species. A natural ecological definition of output power is the energy flux through top predator or consumer species (Saavedra et al. (2025); Odum & Pinkerton (1955)). Because this flux can integrate contributions from multiple species and trophic levels, it is natural to ask whether maximizing predator gain also expands the conditions under which the full community can coexist.

Here, we address this question by comparing two quantities defined at different levels of ecological organization. At the level of a focal consumer, we quantify energetic gain through predator output power. At the level of the community, we quantify collective persistence through the probability of feasibility, that is, the range of growth conditions compatible with positive coexistence. We ask whether maximizing output power with respect to interspecific encounter rates also maximizes the probability of feasibility, and thus whether local energetic advantage translates into collective benefit.

Across a progression of food-web structures, we show that the answer is determined by architecture. In the simplest two-species predator–prey system, predator advantage and collective persistence do not align universally. In trophic chains, by contrast, stronger trophic interactions make them mutually reinforcing. Omnivory reverses this pattern. In particular, when omnivory involves the basal trophic level, the interaction strengths that maximize predator output power are shifted toward the boundary of coexistence, where the intermediate consumer is lost. Omnivory is therefore not the starting assumption of this study, but the structural condition that reveals a more general principle: food-web architecture determines whether optimization at one level of organization scales up as collective persistence or instead produces a coexistence cost. We discuss this result in light of evidence that herbivores face a higher extinction risk than other trophic groups (Atwood et al. (2020)), suggesting that conflicts between local gain and collective persistence may be ecologically consequential.

## Methods

Our objective is to compare two quantities defined at different levels of ecological organization. At the level of a focal species, we quantify energetic gain through the output power of the top predator. At the level of the community, we quantify collective persistence through the probability of feasibility. We then ask how these two quantities vary with encounter rates, and whether the interaction strengths that maximize predator gain also maximize the set of conditions under which all species coexist.

### Energy-flux formulation of trophic dynamics

We consider a Lotka–Volterra-type model in which trophic interactions are written explicitly in terms of energy fluxes through species biomasses. The model describes how energy entering at the basal level is redistributed through the trophic network while accounting for maintenance costs, self-regulation, and losses to consumers. For basal species, biomass dynamics reflect energetic input, maintenance losses, density-dependent regulation, and consumption by higher trophic levels. The equations for basal species are

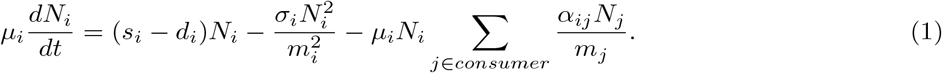

For non-basal species, biomass dynamics reflect maintenance losses, self-regulation, energetic gains from resource consumption, and losses to higher-level consumers. The equations for species at higher trophic levels are

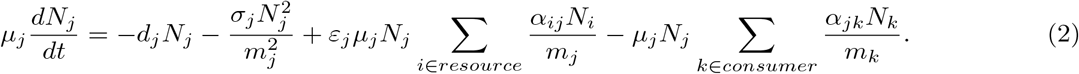

Here, *N*_*i*_ denotes the biomass of species *i* ∈ *S*, where *S* is the species pool. The parameters *µ*_*i*_, *s*_*i*_, and *d*_*i*_ are the mass-specific rates of energy conversion, uptake, and maintenance, respectively. The parameter *σ*_*i*_ denotes the rate of energy exchange within species. Encounter rates are given by *α*_*ij*_ for each consumer– resource pair (*i, j*), and *ε*_*j*_ denotes the assimilation efficiency of consumer species *j*. In this formulation, encounter rates are the key interaction parameters: they govern both the transfer of energy through the network and the coexistence properties of the resulting equilibrium states. Our analysis is therefore organized around how energetic gain and collective persistence change as functions of these encounter rates.

### Output power

To quantify predator-level energetic gain, we define output power as the energy flux through the top consumer. For a top consumer *n* ∈ *S*, the corresponding output power is

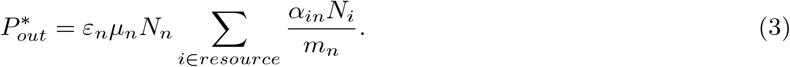

This quantity measures the total equilibrium energy flux entering the focal predator from its resources and therefore provides a natural measure of predator-level energetic gain for a given set of encounter rates. To evaluate whether local energetic gain aligns with collective persistence, we optimize both 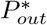 and the probability of feasibility Ω (see Box 1 for details) with respect to encounter rates and compare the locations of their optima. We also consider an upper bound *Q* on the energy supply rate to the system, such that the basal uptake rate satisfies

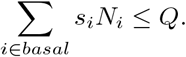

This energetic constraint allows us to test whether the relationship between predator gain and collective persistence changes when total energy input into the system is limited.

### Analytical strategy across network structures

To facilitate analysis, we first consider a simplified version of equations 1 and 2 with homogeneous encounter rates, *α*_*ij*_ = *α* for all consumer–resource pairs. We also define

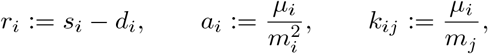

where *k*_*ij*_ is defined for each resource–consumer pair (*i, j*). These simplifications allow us to derive analytical expressions for many quantities of interest. The case of heterogeneous encounter rates is treated in Supplementary Information section S7.

We proceed from simple to more structured trophic networks. We begin with the two-species predator–prey system, which provides the minimal setting in which to compare predator output power and probability of feasibility. This case establishes whether the two objectives align even in the simplest interaction motif. We then turn to a three-species trophic chain to determine whether trophic organization can generate such alignment. Finally, we analyze three-species systems with omnivory, in which the top predator feeds across trophic levels. This progression allows us to isolate the structural condition under which maximizing predator gain ceases to support collective persistence. In the Supplementary Information, we extend the same logic to systems with more trophic levels, different network structures, heterogeneous encounter rates, and type II functional responses. These extensions allow us to assess whether the tension identified in the minimal omnivory motif persists in more general settings.

## Results

We organize the Results around a simple question: when do the interaction strengths that maximize predator advantage also support collective persistence? We begin with the minimal two-species system to establish a baseline, then show that trophic chains can generate alignment between local energetic advantage and coexistence, and finally identify omnivory as the structural condition under which this alignment breaks down. This progression distinguishes what is generic from what is imposed by food-web structure. We first consider the two-species predator–prey system. This case establishes whether any alignment between species-level energetic gain and community-level persistence is already present in the simplest ecological interaction, or whether it instead emerges only in more structured food webs.

### Minimal predator–prey systems provide no generic alignment between predator advantage and collective persistence

For the two-species system, many quantities of interest can be computed analytically. Under the simplifying assumption *µ*_1_ = *µ*_2_ = *σ*_1_ = *σ*_2_ = 1, both the encounter rate that minimizes the probability of feasibility and the encounter rate *α*_*pow*_ that maximizes output power can be written explicitly. The corresponding expressions are given in Supplementary Equations S1 and S2. The two optima depend on different features of the system. Whereas the minimum of Ω(*α*) is determined entirely by interaction parameters, *α*_*pow*_ also depends on the ratio *ρ* = *r*_1_*/d*_2_ between basal growth and predator maintenance. The values of *α* that maximize output power and feasibility therefore do not generically coincide. Even in the simplest predator–prey system, there is no universal reason to expect that increasing predator gain will enlarge the set of growth conditions compatible with coexistence. This baseline result shows that any systematic alignment between predator advantage and collective persistence must arise from food-web structure rather than from a generic property of predator–prey dynamics. In the Supplementary Information, we also show how the distance between these optima varies with *ρ* (Fig. S1).

### Trophic chains create alignment between predator advantage and collective persistence

Next, we consider a three-species trophic chain consisting of a basal resource, an intermediate consumer, and a top predator. This case asks whether a minimal trophic organization can generate the alignment absent in the two-species system. We first determine *α*_*min*_, the minimum value of *α* for which a feasible solution exists, by imposing *C*\* *>* 0 and *P*\* *>* 0, where *C*\* and *P*\* are the equilibrium biomasses of the intermediate consumer and top predator. The closed-form expression for *α*_*min*_ is given in Supplementary Equation S3. We then compare how the probability of feasibility Ω and output power 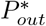 vary with *α*. In three dimensions, Ω can be computed geometrically from the spherical triangle defined by the generators of the feasibility cone, yielding the expression in Supplementary Equation S4. We define output power as the energy flux through the top predator at equilibrium:

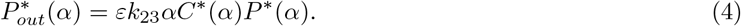

For the parameter values shown in Figure 1, both 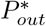 and Ω(*α*) increase monotonically with *α*. This qualitative relationship is unchanged for other parameter values, including the case in which the parameters of the top and basal species are swapped (Fig. S2). Thus, the trophic chain produces a clear alignment between local energetic advantage and collective persistence: stronger trophic interactions simultaneously increase predator gain and enlarge the set of growth conditions compatible with coexistence. This conclusion remains unchanged when energy supply is capped. If *E*(*α*) = *r*_*i*_*R*\*(*α*) ≤ *Q*, the constraint limits the equilibrium biomass of the basal species. Because *R*\*(*α*) decreases monotonically with *α* (Figure 1C), the energy cap restricts the admissible range of interaction strengths without altering the qualitative alignment between the two objectives. Within the permitted range, large values of *α* still maximize both 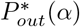 and Ω(*α*). The alignment between predator advantage and collective persistence is therefore not generic, but it can emerge from trophic organization.

**Figure 1:**
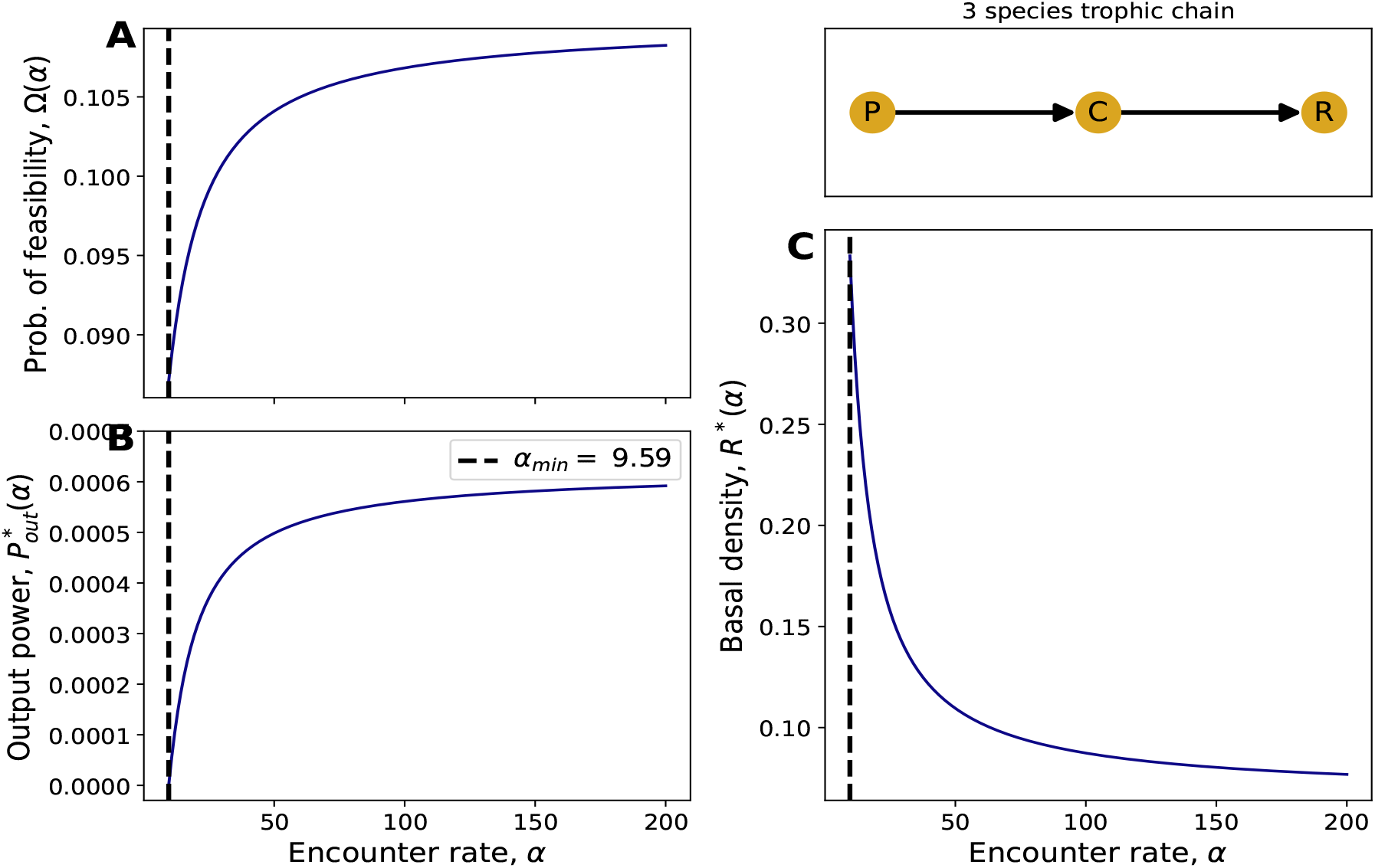
The optima of output power and probability of feasibility match exactly in a 3-species trophic chain. For the case of a 3 species trophic chain, the three subplots A, B and C show the probability of feasibility Ω(*α*), output power 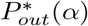 and the equilibrium density of the basal species *R*\*(*α*) respectively. The parameter values are *a*_1_ = 1, *a*_2_ = 4, *a*_3_ = 9, *ε* = 0.1, *k*_12_ = 2, *k*_23_ = 3, *r*_1_ = 1, *d*_2_ = 0.5 and *d*_3_ = 0.1.

### Omnivory decouples predator advantage from collective persistence

We now turn to the central structural case of this study: omnivory involving the basal trophic level. In this motif, the top predator feeds both on the intermediate consumer and directly on the basal species. This added pathway changes the relationship between predator gain and collective persistence because the predator no longer depends exclusively on the persistence of the intermediate trophic level. Denoting the equilibrium biomasses by *N*\* = (*R*\*, *C*\*, *P*\*)^⊤^, we write *N*\* = num(*N*\*)*/D*(*α*). Details of the analytical expressions are given in Supplementary Information section S4. In this case, output power includes a direct contribution from the basal species:

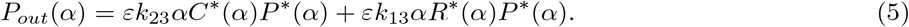

To characterize *α*_*pow*_, we consider two cases depending on whether the feasible region lies to the left or to the right of the root of *D*(*α*), which we denote by *α*_*Dzero*_. In both cases, we choose *a*_1_ = 1, *a*_2_ = 4, *a*_3_ = 9, *k*_12_ = 2, *k*_23_ = *k*_13_ = 3, and *ε* = 0.1. Because *D*(*α*) is independent of the vector ***r*** = (*r*_1_, −*d*_2_, −*d*_3_), varying ***r*** shifts the feasible region relative to *α*_*Dzero*_. In the Supplementary Information, we also show the corresponding results for *a*_1_ = 9, *a*_2_ = 4, *a*_3_ = 1, *k*_12_ = 2, *k*_23_ = *k*_13_ = 1, and *ε* = 0.1, corresponding to the case in which the masses of the top and basal trophic levels are swapped. The qualitative outcome remains unchanged (Fig. S3).

We first choose ***r*** such that feasible solutions exist and *α*_*pow*_ *< α*_*Dzero*_. For *r*_1_ = 1, *d*_2_ = 0.1, and *d*_3_ = 0.3, the finite root of *P*\*(*α*) determines *α*_*min*_, whereas the root of *C*\*(*α*) is the largest value of *α* for which a feasible solution exists; we denote this value by *α*_*max*_ (Figure 2B). Because output power increases monotonically throughout the feasible region, *α*_*pow*_ = *α*_*max*_ (Figure 2B). Thus, the power optimum coincides with the point at which the intermediate consumer lies at the edge of extinction. Supplementary Information section S5 further shows how *α*_*pow*_ varies across ***r***-space.

**Figure 2:**
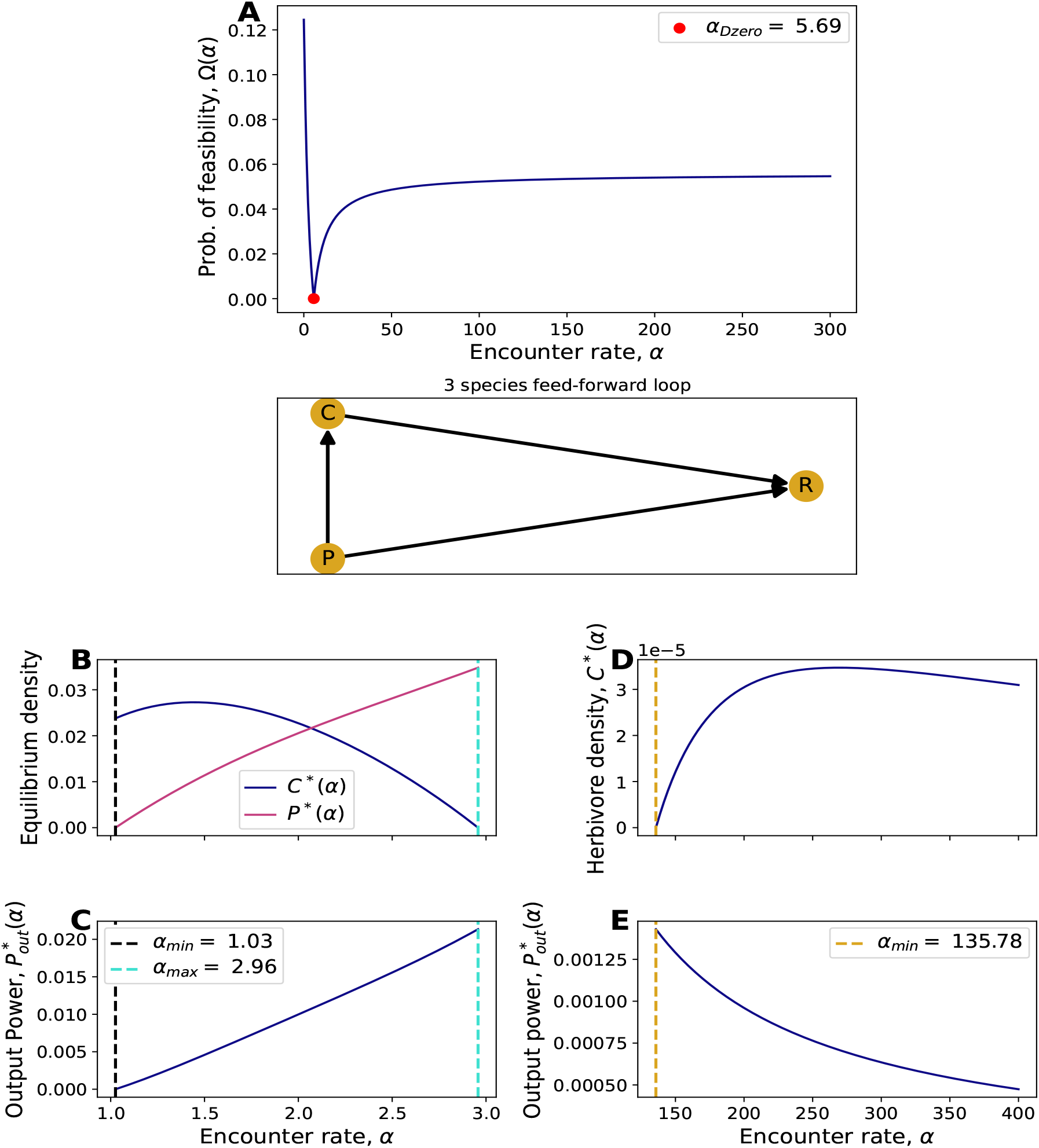
Basal omnivory shows no generic alignment between output power and probability of feasibility. The three species omnivory motif illustrated here is categorized as a feed-forward loop in network theory literature Milo et al. (2002). All the panels show results for parameter values *a*_1_ = 1, *a*_2_ = 4, *a*_3_ = 9, *ε* = 0.1, *k*_12_ = 2, *k*_23_ = 3 and *k*_13_ = 3. Panel **A** shows the probability of feasibility as a function of *α*, which is independent of the vector ***r***. We also highlight two different ways in which 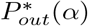 varies with *α*. Panels **B** and **C** correspond to ***r*** = (*r*_1_, −*d*_2_, −*d*_3_) = (1, −0.1, −0.3), while panels **D** and **E** have ***r*** = (0.6, −0.1, −1).

We then consider the complementary case, with ***r*** = (0.6, −0.1, −1) and all other parameters unchanged. Here, the larger predator maintenance rate shifts the feasible region to higher values of *α*, and with it the location of *α*_*pow*_ (Figure 2E). In this regime, the root of *C*\*(*α*) determines *α*_*min*_ (Figure 2D). This root also coincides with *α*_*pow*_, because output power decreases monotonically for *α > α*_*min*_. Maximizing output power therefore again pushes the intermediate trophic level to extinction. At the same time, feasibility increases as one moves away from *α*_*pow*_ (Figures 2A and 2E), showing that the power optimum lies at the boundary of coexistence rather than in the region where coexistence is most broadly supported.

Although the position of *α*_*pow*_ relative to *α*_*Dzero*_ depends on the growth-rate vector, the qualitative outcome is the same in both cases: maximizing predator gain pushes the system toward the coexistence boundary, marked here by the loss of the intermediate trophic species. This pattern is not restricted to the minimal three-species motif. The extinction of the herbivore at *α*_*pow*_ remains the norm whenever omnivory involves the basal species. We show that this remains true with additional trophic levels (Fig. S8), a type II functional response (Fig. S13), and heterogeneous encounter rates (Fig. S7). The optima of output power and feasibility are also misaligned in cases with non-basal omnivory, but the specific pattern of herbivore extinction characterizes only networks that exhibit basal omnivory alone (Figs. S9 and S10). Omnivory therefore reveals a structural tension between predator advantage and collective persistence: the interaction strengths that maximize predator gain are no longer those that maximize the range of conditions under which all species persist.

## Discussion

Our study addresses a broad question in biological organization: when does local advantage scale up as a benefit to the whole system, and when does it instead generate a collective cost? In food webs, we show that the answer depends on architecture. The interaction strengths that maximize predator output power do not generically coincide with those that maximize the probability of feasibility. In the simplest two-species system, there is no universal alignment between the two objectives. In trophic chains, by contrast, predator advantage and collective persistence can align. Basal omnivory reverses this pattern: when the top predator also feeds directly on the basal resource, the power optimum is shifted towards the boundary of coexistence, where the intermediate trophic level is lost.

The mechanism is straightforward. In a trophic chain, energy reaches the top predator only through the intermediate consumer, so increasing interaction strength can enhance predator gain while preserving the integrity of the chain. Under basal omnivory, however, the predator gains direct access to the basal resource. This alternative energetic pathway relaxes the requirement that the intermediate species persist, with the result that the interaction strengths maximizing predator gain are shifted towards states in which coexistence breaks down. Basal omnivory therefore decouples predator performance from community persistence.

This result refines how omnivory should be interpreted in the broader food-web literature. Because omnivory increases connectance, and therefore complexity when species richness is fixed, it was initially thought to be destabilizing and rare in natural systems (Pimm & Lawton (1978)). Later work showed that omnivory is widespread (Thompson et al. (2007)), and that it can promote stability when feeding is distributed across many weak interactions and trophic levels (McCann & Hastings (1997); Kratina et al. (2012)). Our results address a different question: not whether omnivory can stabilize communities in general, but whether the interaction strengths that maximize predator gain are also those most compatible with coexistence. From this perspective, basal omnivory is distinctive because it places the power optimum at, or very near, the coexistence boundary (Figures 2, S3, S13 and S7). This conclusion remains unchanged with additional trophic levels, heterogeneous encounter rates, and type II functional responses. By contrast, non-basal omnivory also creates tension between power and feasibility, but does not produce the same systematic extinction of the intermediate consumer (Figures S9 and S10).

These findings also recast the maximum power principle through the lens of collective persistence. In the presence of basal omnivory, maximum-power states correspond to low feasibility and occur precisely where the herbivore goes extinct. Thus, even if real ecosystems are observed in nearby sub-maximal states, relatively small shifts in interaction strengths could move them towards herbivore loss. This interpretation is also consistent with theory on intraguild predation, in which two consumers share a resource while one also preys on the other (Polis et al. (1989); Arim & Marquet (2004)). Theory has shown that coexistence in such systems requires the intermediate species to remain the superior competitor for the shared resource, while the predator must benefit sufficiently from consuming that intermediate species (Holt & Polis (1997)). We find that these conditions fail at the maximum-output-power threshold, where the predator can sustain its energetic gain through the basal resource alone and the herbivore is lost. The ecological implications may be especially relevant when food webs are perturbed. Experimental work shows that invasive omnivorous fish can reduce the density and temporal stability of herbivorous fish while leaving basal resources relatively unaffected (Long et al. (2011)). More broadly, herbivores have been found to face higher extinction risk than carnivores or omnivores across major vertebrate groups worldwide (Atwood et al. (2020)). Our results suggest one possible mechanism for this asymmetry: when omnivorous predators gain direct energetic access to basal resources, the states that maximize predator gain need not support persistence of the herbivore.

By revisiting a long-standing energetic principle through a systematic analysis of feasibility, we provide theoretical expectations for when local energetic advantage and collective persistence align, and when they diverge. Basal omnivory is the key structural condition that reveals this tension most clearly. More broadly, our results show that food-web architecture governs whether optimization at one level of organization scales up as a collective benefit or instead generates a coexistence cost. Future experiments could test these predictions in simple systems exhibiting intraguild predation. More generally, we advocate wider use of ecological models that treat energy as a central currency, because such approaches can reveal when local energetic gains support collective persistence, and when they undermine it.

## Acknowledgements

The authors would like to thank Chengyi Long for useful discussions. A.V. is thankful to the Wallenberg Foundation for funding. S.S. acknowledges support from the National Science Foundation under Grant No. DEB-2436069.

## Data accessibility statement

No new data were used in the preparation of this manuscript.

## Code availability

The associated code is available at this github repository.

## Box 1

Quantifying collective persistence and predator advantage

We compare two quantities that capture different aspects of food-web organization. The first is predator output power, defined as the equilibrium energy flux through the top consumer. It measures the local energetic advantage achieved by the predator for a given set of encounter rates. The second is the probability of feasibility, denoted by Ω, which measures the fraction of growth conditions under which all species coexist at positive equilibrium.

To compute Ω, we use the feasibility domain of the interaction network. This domain is the set of intrinsic growth-rate vectors that yield positive coexistence equilibria for a given interaction matrix. Geometrically, it is a cone generated by the columns of the negative interaction matrix. Its solid angle quantifies the fraction of growth conditions compatible with coexistence, and therefore determines the value of Ω.

Comparing predator output power and Ω as functions of encounter rates allows us to ask whether local energetic advantage and collective persistence are aligned. When the same encounter rates increase both quantities, predator advantage reinforces coexistence. When the encounter rates that maximize output power instead lie near the boundary of the feasibility domain, local energetic gain comes at a collective

## Supplementary Information

### S1 Interaction matrices and reduced formulation

To facilitate analytical progress, we introduce the reduced quantities

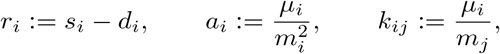

and write the generalized Lotka–Volterra system as

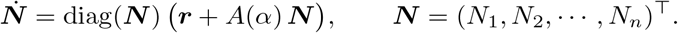

For trophic-chain systems with a single encounter-rate parameter *α*, the reduced interaction matrix takes the form

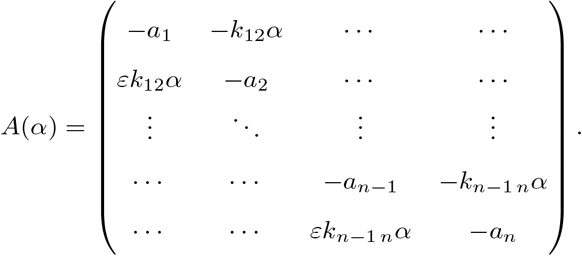

The corresponding feasibility domain is the simplicial cone generated by the columns of −*A*(*α*). In the main text, we quantify the breadth of this cone by its normalized solid angle.

### S2 Two-species system: analytical optima

For the two-species system, many quantities of interest can be computed analytically. Assuming *µ*_1_ = *µ*_2_ =*σ*_1_ = *σ*_2_ = 1, the value of *α* at which the probability of feasibility is minimized is

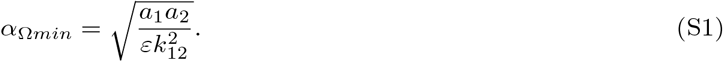

The output power varies unimodally with *α*. If *α*_*pow*_ denotes the encounter rate that maximizes output power, then

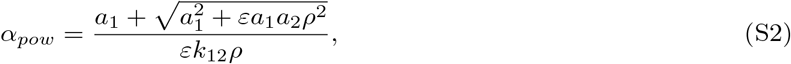

where

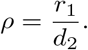

Figure 1 shows how the distance between *α*_Ω*min*_ and *α*_*pow*_ varies with *ρ*.

**Figure S1:**
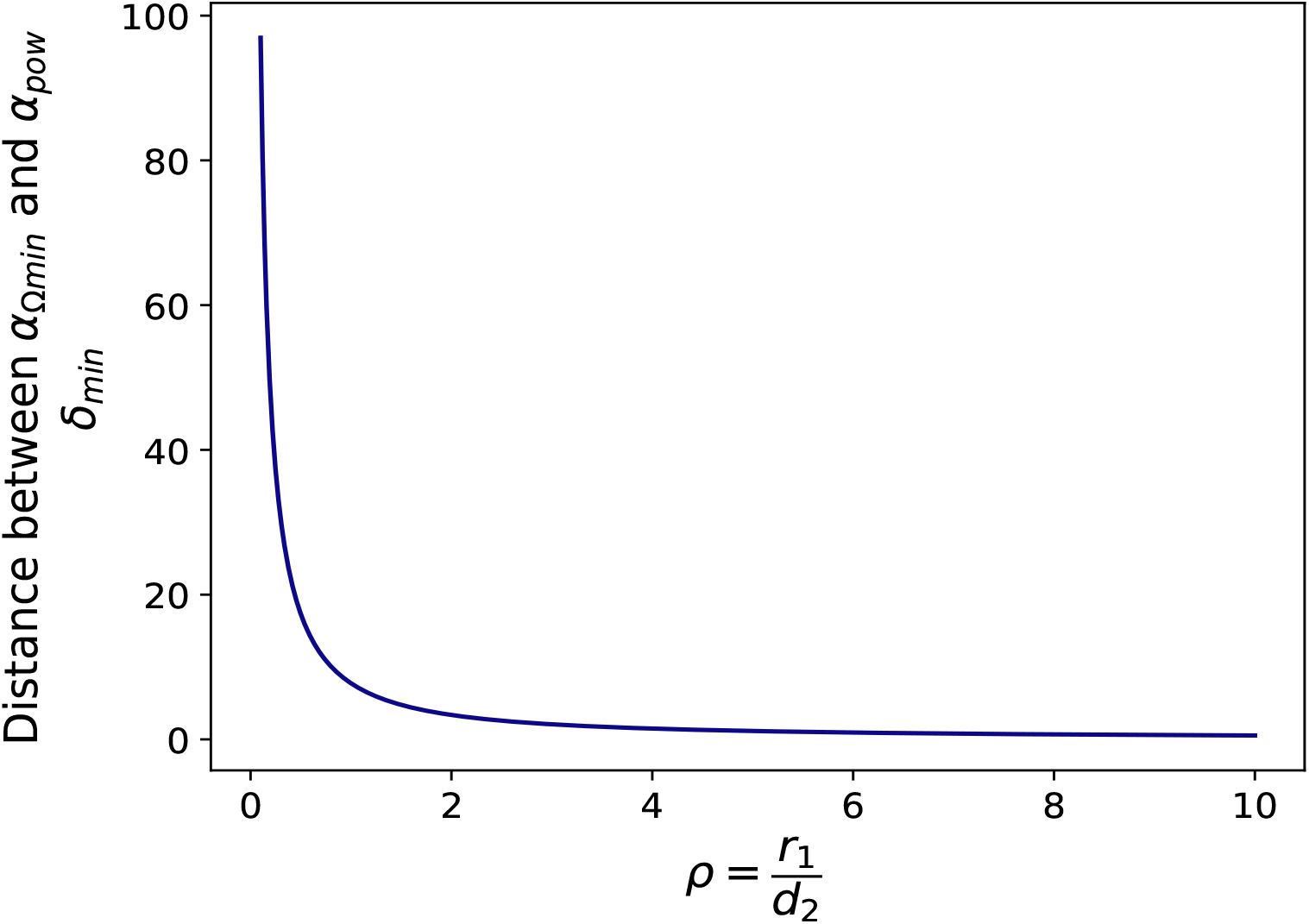
Distance between the encounter rate that minimizes the probability of feasibility and the encounter rate that maximizes output power in the two-species system, plotted as a function of *ρ* = *r*_1_*/d*_2_.

### S3 Three-species trophic chain

For the three-species trophic chain, the minimum encounter rate required for a feasible solution is determined by the condition *C*\* *>* 0:

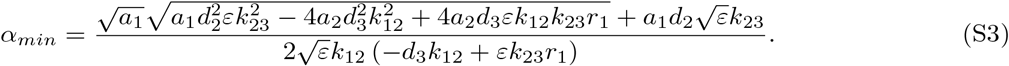

In three dimensions, the probability of feasibility can be computed from the spherical triangle spanned by the normalized generators 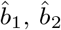 and 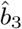of the feasibility cone:

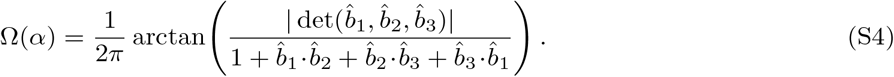

These expressions underlie the monotonic increase of both output power and probability of feasibility shown in Figure 1.

**Figure S2:**
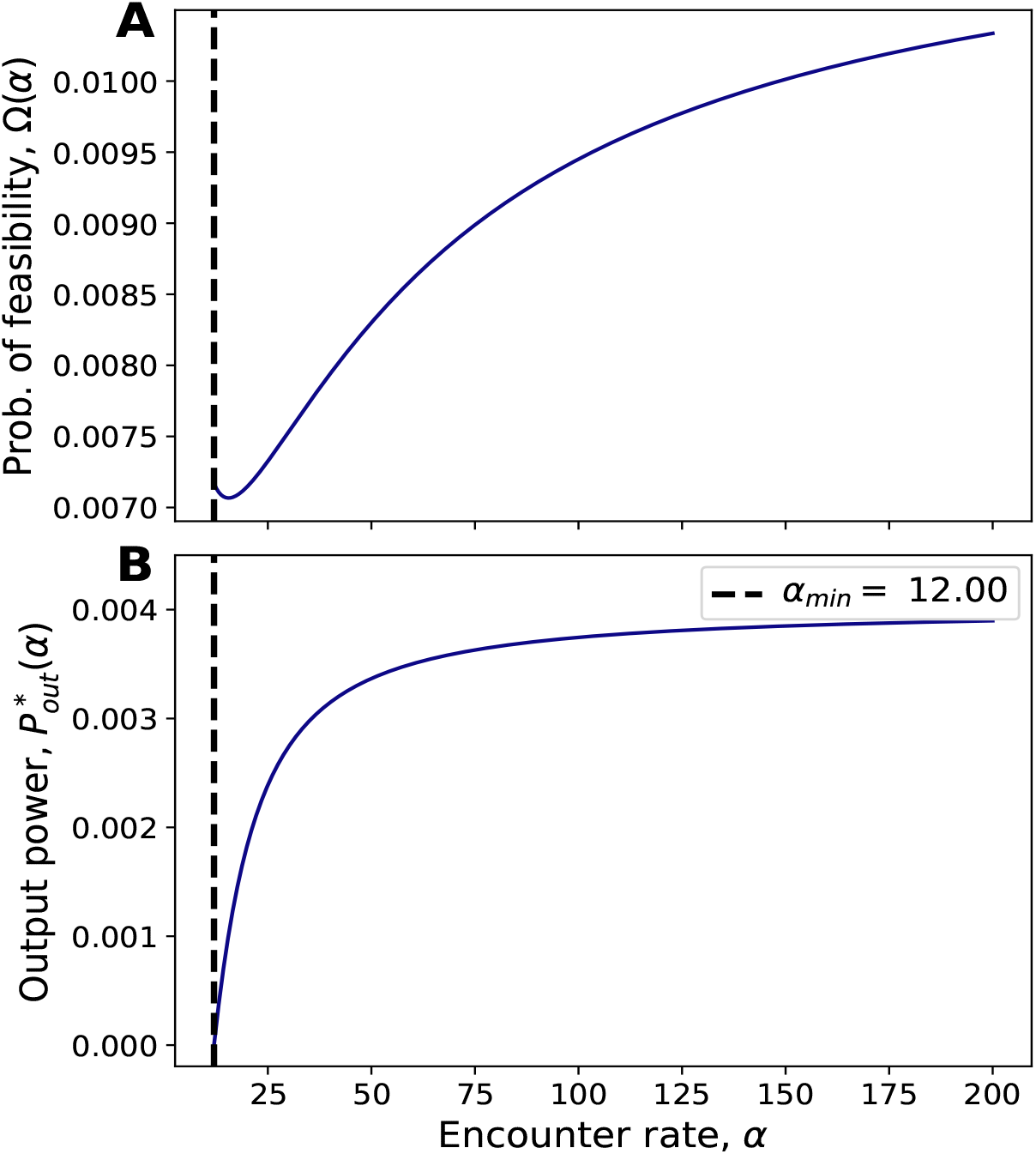
Probability of feasibility Ω(*α*) and output power 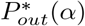 for the three-species trophic chain when the parameters of the top and basal species are swapped. Parameter values are *a*_1_ = 9, *a*_2_ = 4, *a*_3_ = 1, *ε* = 0.1, *k*_12_ = 2, *k*_23_ = 1, *r*_1_ = 4, *d*_2_ = 0.2 and *d*_3_ = 0.1.

### S4 Three-species omnivory: equilibrium expressions

When the top predator also feeds on the basal species, the interaction matrix becomes

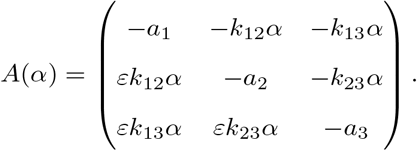

Writing the equilibrium biomasses as *N*\* = (*R*\*, *C*\*, *P*\*)^⊤^= *num*(*N*\*)*/D*(*α*), the numerators are

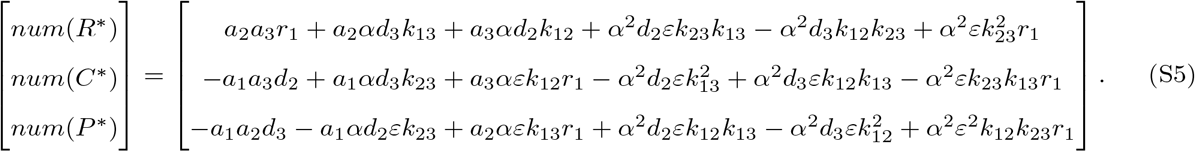

The common denominator is

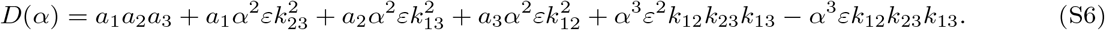

Because *num*(*R*\*), *num*(*C*\*) and *num*(*P*\*) are quadratic in *α*, whereas *D*(*α*) is cubic, the equilibrium biomasses can display sharp changes near roots of *D*(*α*). The location of *α*_*pow*_ relative to these roots determines the two cases described in the main text.

**Figure S3:**
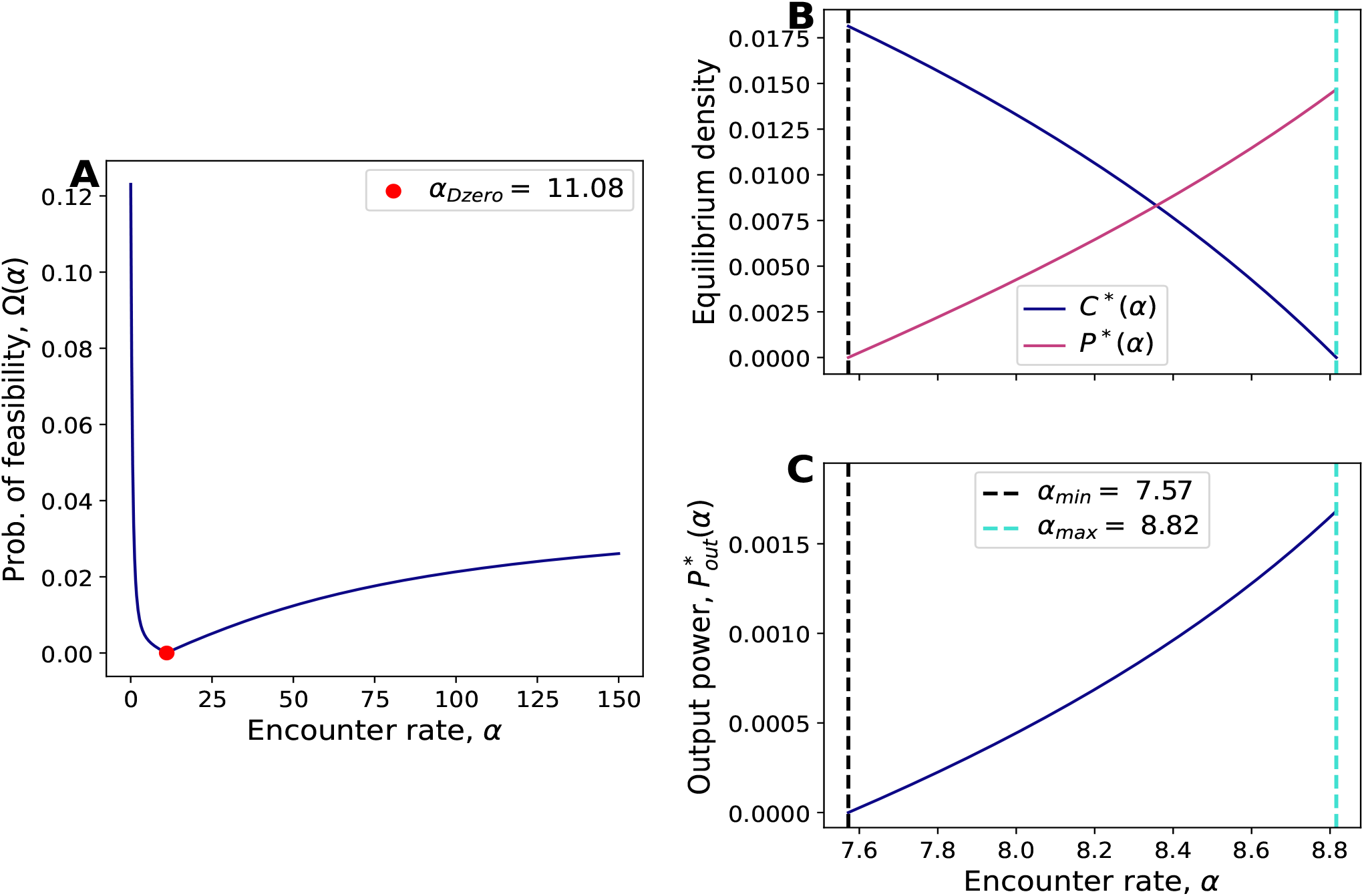
Three-species basal-omnivory case with the parameters of the top and basal species swapped. Parameter values are *a*_1_ = 9, *a*_2_ = 4, *a*_3_ = 1, *ε* = 0.1, *k*_12_ = 2, *k*_23_ = 1, *k*_13_ = 1, and ***r*** = (*r*_1_, −*d*_2_, −*d*_3_) = (1.3, −0.1, −0.1).

### S5 *α*_*pow*_ in *r*-space

To explore how the power optimum changes across growth-rate vectors, we rewrite the relevant ratios as

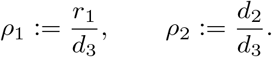

It is then useful to compare *α*_*pow*_ with the point of minimum feasibility, which in the basal-omnivory case coincides with *α*_*Dzero*_. We therefore define

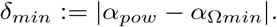

Figure 4 shows how *δ*_*min*_ varies across (*ρ*_1_,*ρ*_2_)-space.

**Figure S4:**
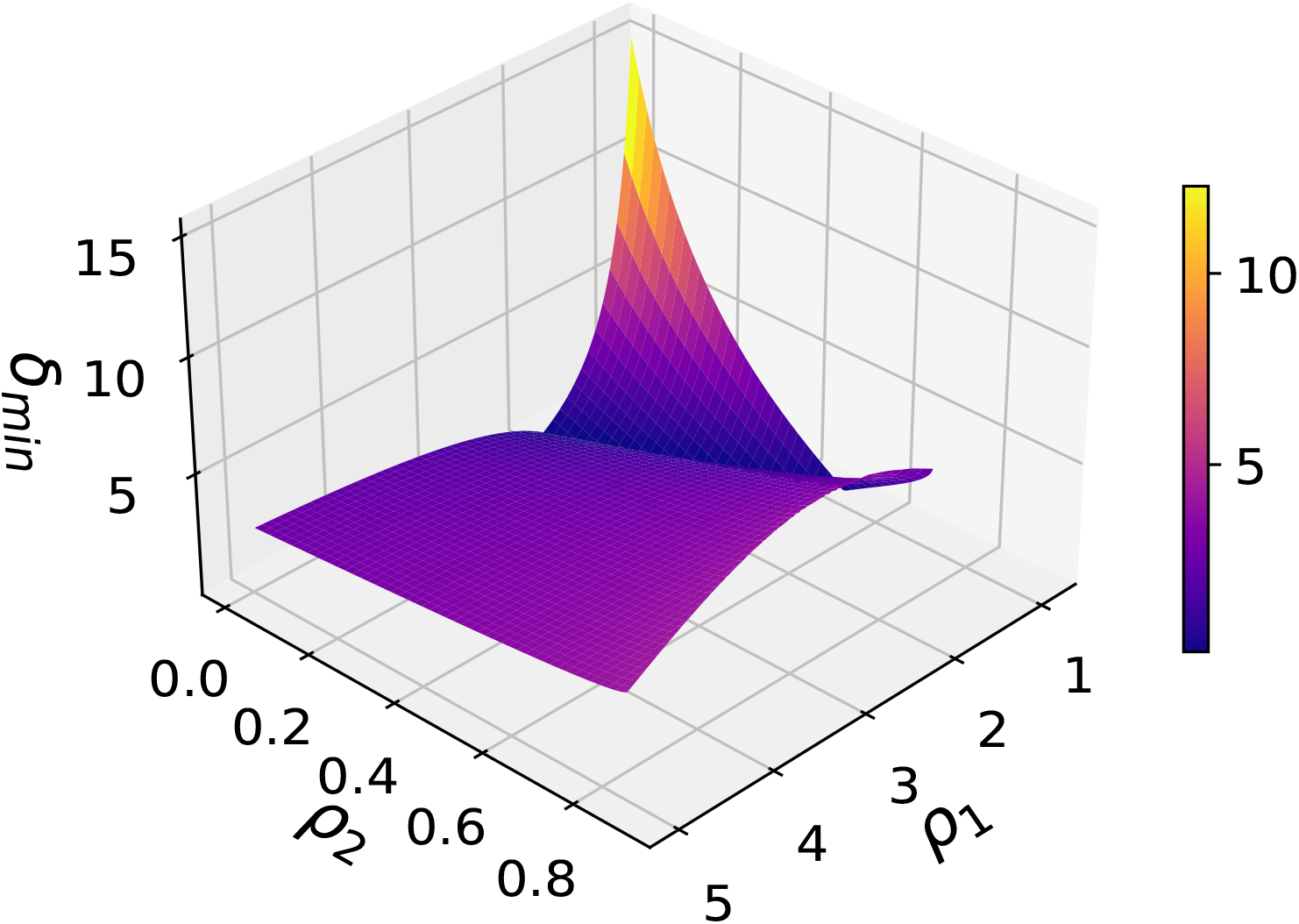
Distance *δ*_*min*_ between the encounter rate that maximizes output power and the encounter rate of minimum feasibility in the three-species basal-omnivory system, shown as a function of *ρ*_1_ = *r*_1_*/d*_3_ and *ρ*_2_ = *d*_2_*/d*_3_. Parameter values are *a*_1_ = 1, *a*_2_ = 4, *a*_3_ = 9, *k*_12_ = 2, *k*_23_ = *k*_13_ = 3, and *ε* = 0.1. Regions without values correspond to growth-rate vectors for which no feasible solution exists.

### S6 Two basal species and one consumer

The network of interactions is slightly changed such that the system can be studied using the following interaction matrix:

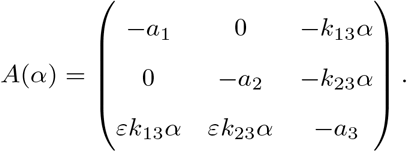

The output power for this system has the following expression:

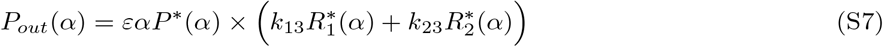

where the equilibrium densities of resource 1, resource 2 and the consumer are denoted by 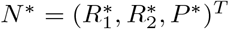. We write down the closed form expressions for these as *N*\* = *num*(*N*\*)*/D*(*α*) where the denominator *D*(*α*) is the same across the three densities.

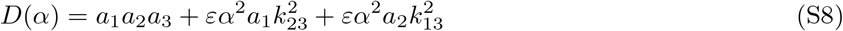

The numerators have the following expressions:

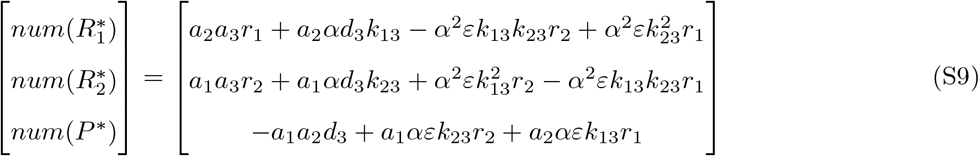

To give a general impression of how output power and feasibility vary with *α*, we work with the following parameter choices: *a*_1_ = 1, *a*_2_ = 4, *a*_3_ = 9, *k*_13_ = *k*_23_ = 3, *ε* = 0.1.

**Figure S5:**
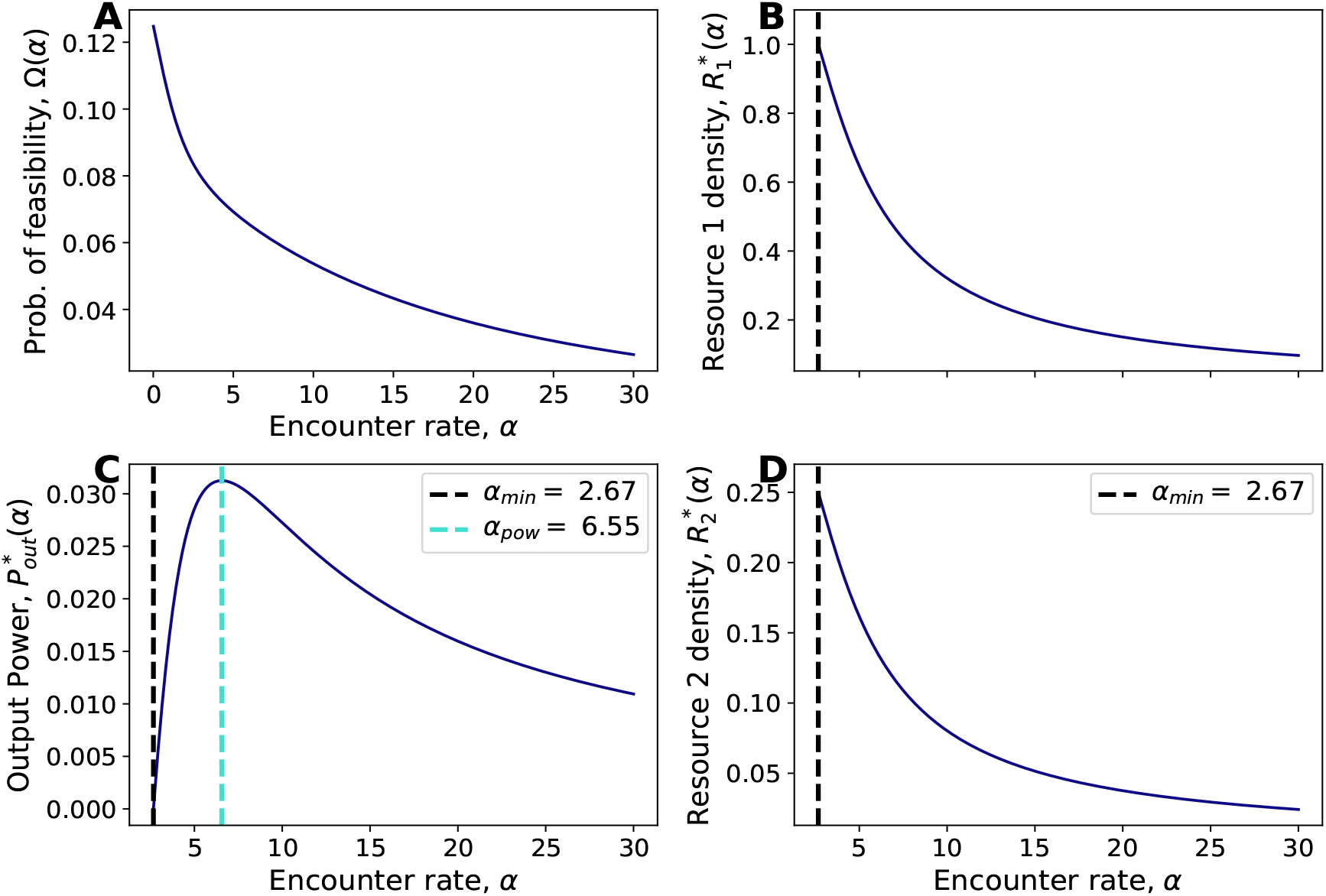
For a food-web with one consumer and two resources, the subplots A and C show the probability of feasibility Ω(*α*) and output power 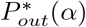 versus encounter rate *α* respectively. In B and D, we plot the resource densities 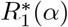 and 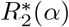. Enforcing a supply cap, i.e. 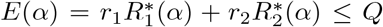 moves *α*_*min*_ to the right, such that both 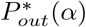 and Ω(*α*) align. Parameter values are *a*_1_ = 1, *a*_2_ = 4, *a*_3_ = 9, *k*_13_ = *k*_23_ = 3, *ε* = 0.1, *r*_1_ = 1, *r*_2_ = 1, and *d*_3_ = 1.

In Figure S5, the probability of feasibility Ω(*α*) and the output power 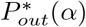 show a mismatch in trends at lower values of *α*. However, if we limit the energy supply to the system such that 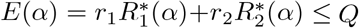, then *α*_*min*_ would shift to higher values of *α* to accommodate the reduced maxima of 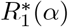 and 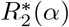 (Figure S5B and S5D). This guarantees the alignment of the two objectives.

Notably, if the parameters for one of the resources are swapped with those of the consumer, then Ω(*α*) and 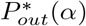 are unimodal in *α* and show perfect alignment in their optima (Figure S6).

**Figure S6:**
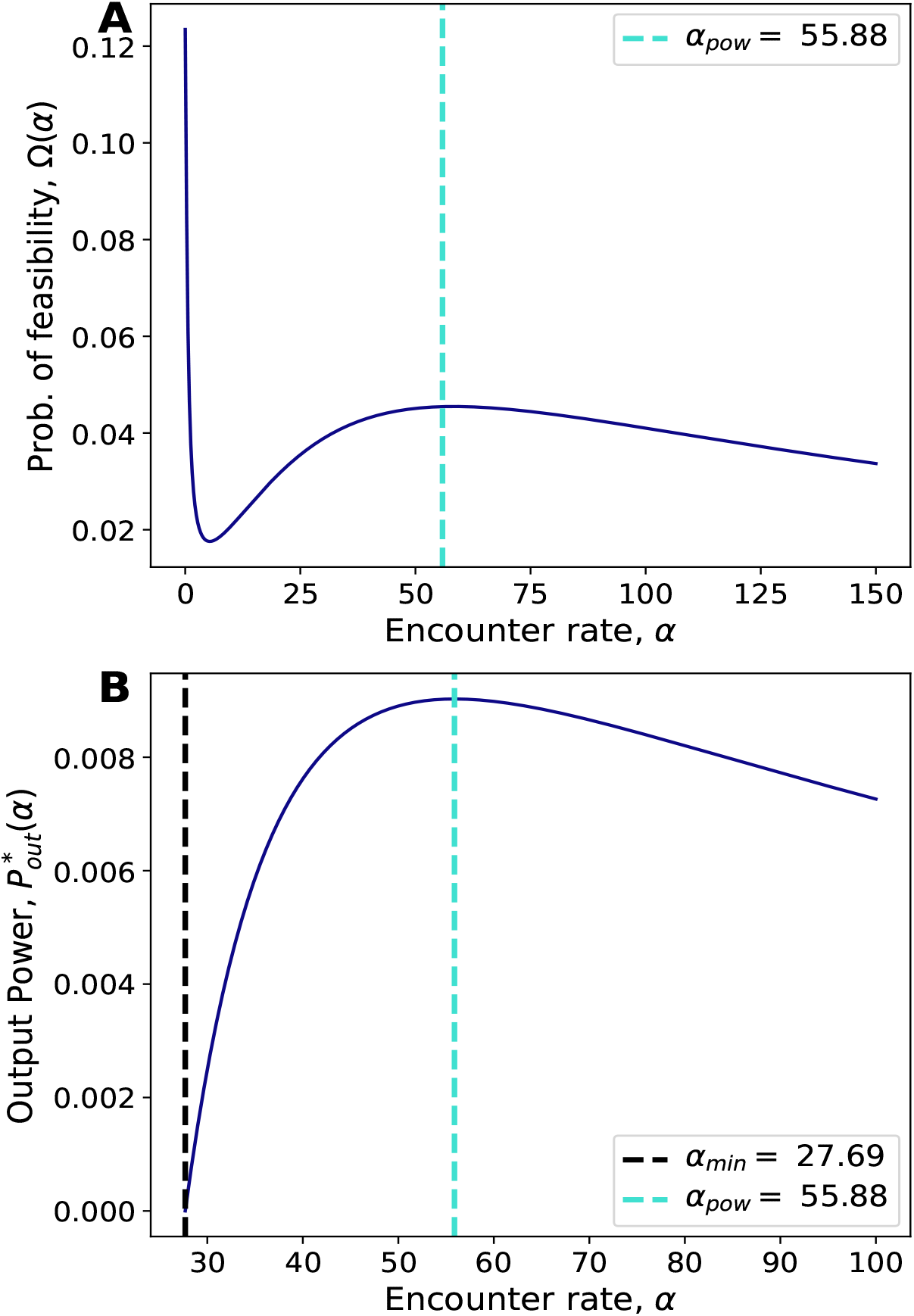
For a food-web with one consumer and two resources, this figure considers the case where the parameters of one of the resources are swapped with those of the consumer. The subplots A and B show the probability of feasibility Ω(*α*) and output power 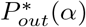 versus encounter rate *α* respectively. Parameter values are *a*_1_ = 9, *a*_2_ = 4, *a*_3_ = 1, *k*_13_ = *k*_23_ = 1, *ε* = 0.1, *r*_1_ = 1, *r*_2_ = 1, and *d*_3_ = 1.

### S7 Heterogeneity in encounter rates

We also consider the case in which the omnivory encounter rate differs from the other encounter rates. Denoting the basal-omnivory encounter rate by *α*_*omni*_, the interaction matrix becomes

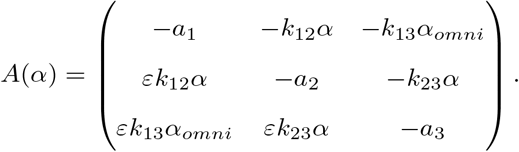

We use the same parameter values as in the corresponding main-text case, with *a*_1_ = 1, *a*_2_ = 4, *a*_3_ = 9, *k*_12_ = 2, *k*_23_ = *k*_13_ = 3, *ε* = 0.1, *r*_1_ = 1, *d*_2_ = 0.1, and *d*_3_ = 0.3. Figure 7 shows that even when encounter rates are allowed to differ, output power is maximized at the point where the intermediate trophic species goes extinct.

**Figure S7:**
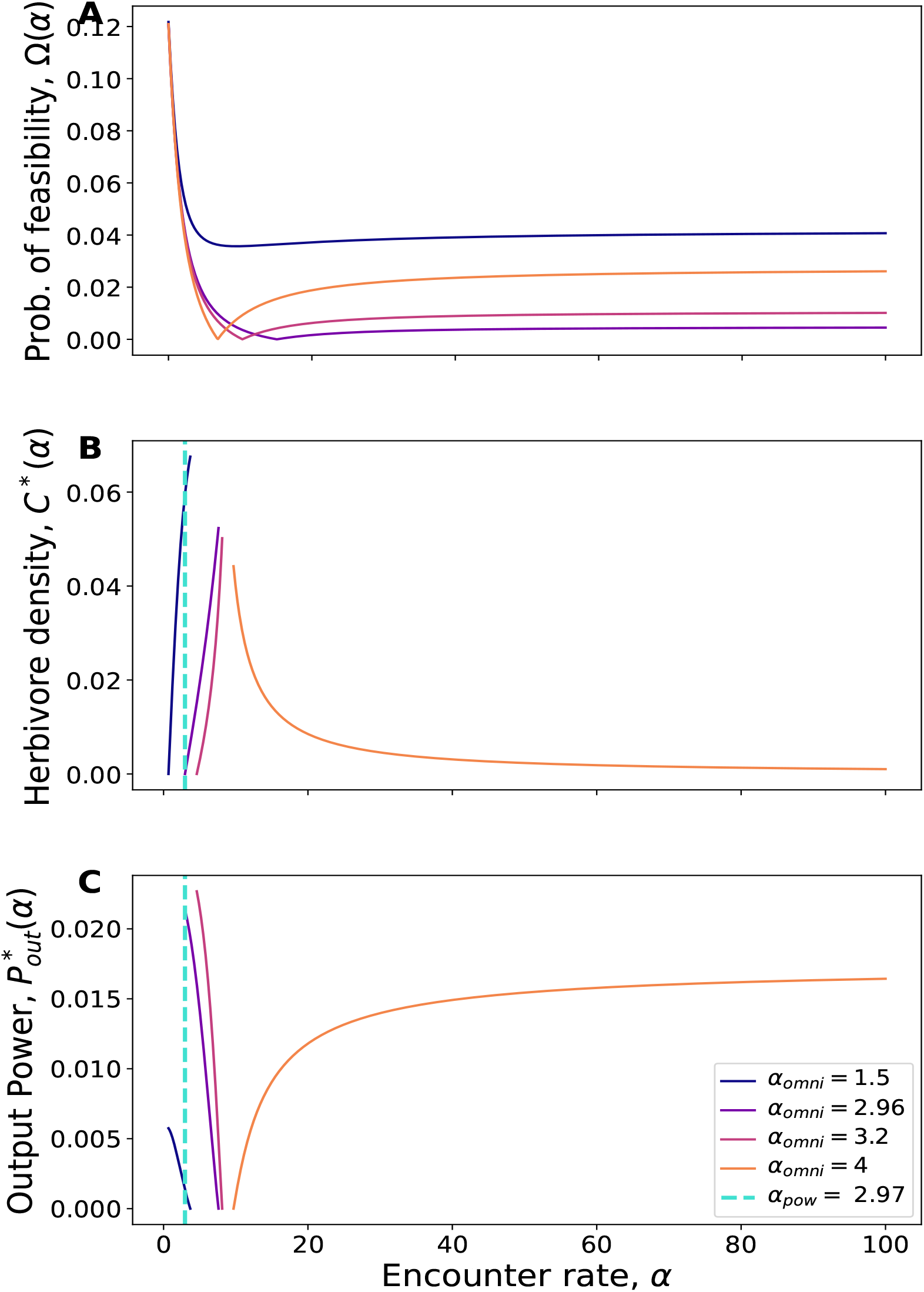
Output power in the feasible region for different values of the omnivory encounter rate *α*_*omni*_. The dashed line highlights that, for *α*_*omni*_ = 2.96, output power is maximized at the same value of *α* for which *C*\*(*α*) goes to zero. Parameter values are *a*_1_ = 1, *a*_2_ = 4, *a*_3_ = 9, *k*_12_ = 2, *k*_23_ = *k*_13_ = 3, *ε* = 0.1, *r*_1_ = 1, *d*_2_ = 0.1, and *d*_3_ = 0.3.

### S8 Basal and non-basal omnivory in four trophic levels

Using four trophic levels, we analyze how the position of omnivory within the network modifies the relationship between output power and coexistence.

We analyze three cases that are characterized by the following interaction matrices:

#### Omnivory without the top trophic level

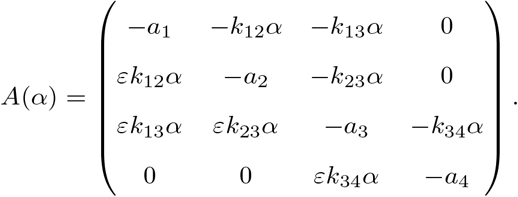

#### Omnivory without the basal species

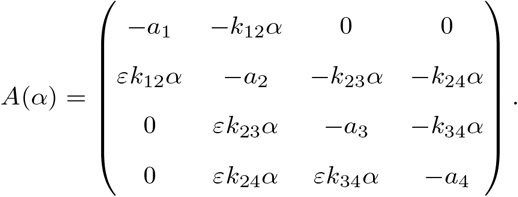

#### Hierarchy of basal and non-omnivory

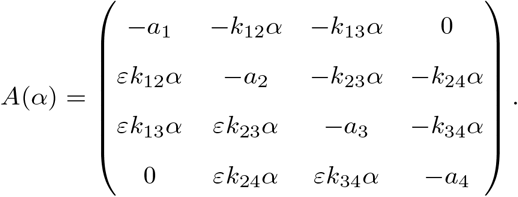

For the case of omnivory without the top predator, the qualitative picture matches that of the three-species basal-omnivory motif: the output-power optimum coincides with the extinction threshold of the herbivore-like intermediate species (Figure S8). By contrast, when omnivory excludes the basal species, output power is no longer tied to the extinction of the intermediate trophic level (Figures S9 and S10).

Even for the case of non-omnivory, the optima of output power and probability of feasibility do not align. While we cannot analytically compute Ω(*α*) for 4 species systems, we can identify the minimum of Ω(*α*) by plotting *D*\*(*α*) which is the denominator of the equilibrium densities. As in figure 2, *α*_*Dzero*_ is the point at which Ω(*α*) is minimum (Figure S11). Therefore, Ω(*α*) would decrease monotonically in *α* till *α*_*Dzero*_, even though 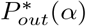 increases in parts of that region (Figure S10).

**Figure S8:**
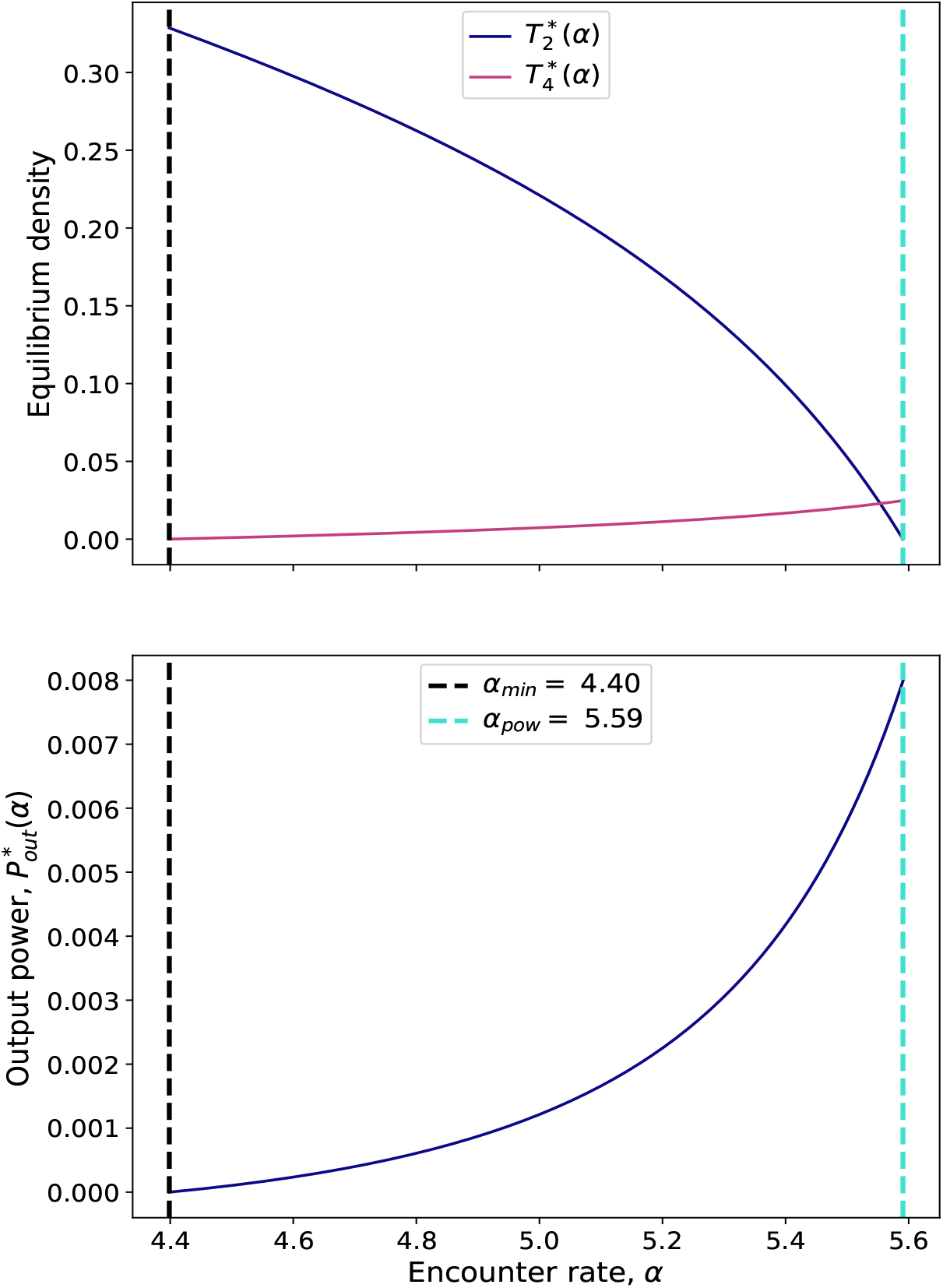
Four-trophic-level system with omnivory below the top predator. As in the three-species basal-omnivory case, the output-power optimum coincides with the extinction of the herbivore-like species 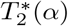. Parameter values are *a*_1_ = 1, *a*_2_ = 2, *a*_3_ = 4, *a*_4_ = 9, *k*_12_ = 1.414, *k*_23_ = 2, *k*_34_ = 3, *k*_13_ = 2, *ε* = 0.1, *r*_1_ = 5, *d*_2_ = 0.1, *d*_3_ = 2, and *d*_4_ = 0.1.

### S9 Type II functional response

We also consider the corresponding systems with saturating type II functional responses. For the trophic chain, the dynamics are

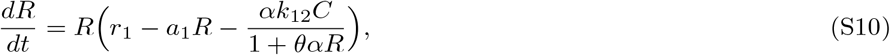

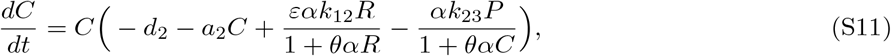

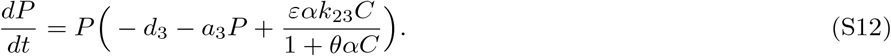

**Figure S9:**
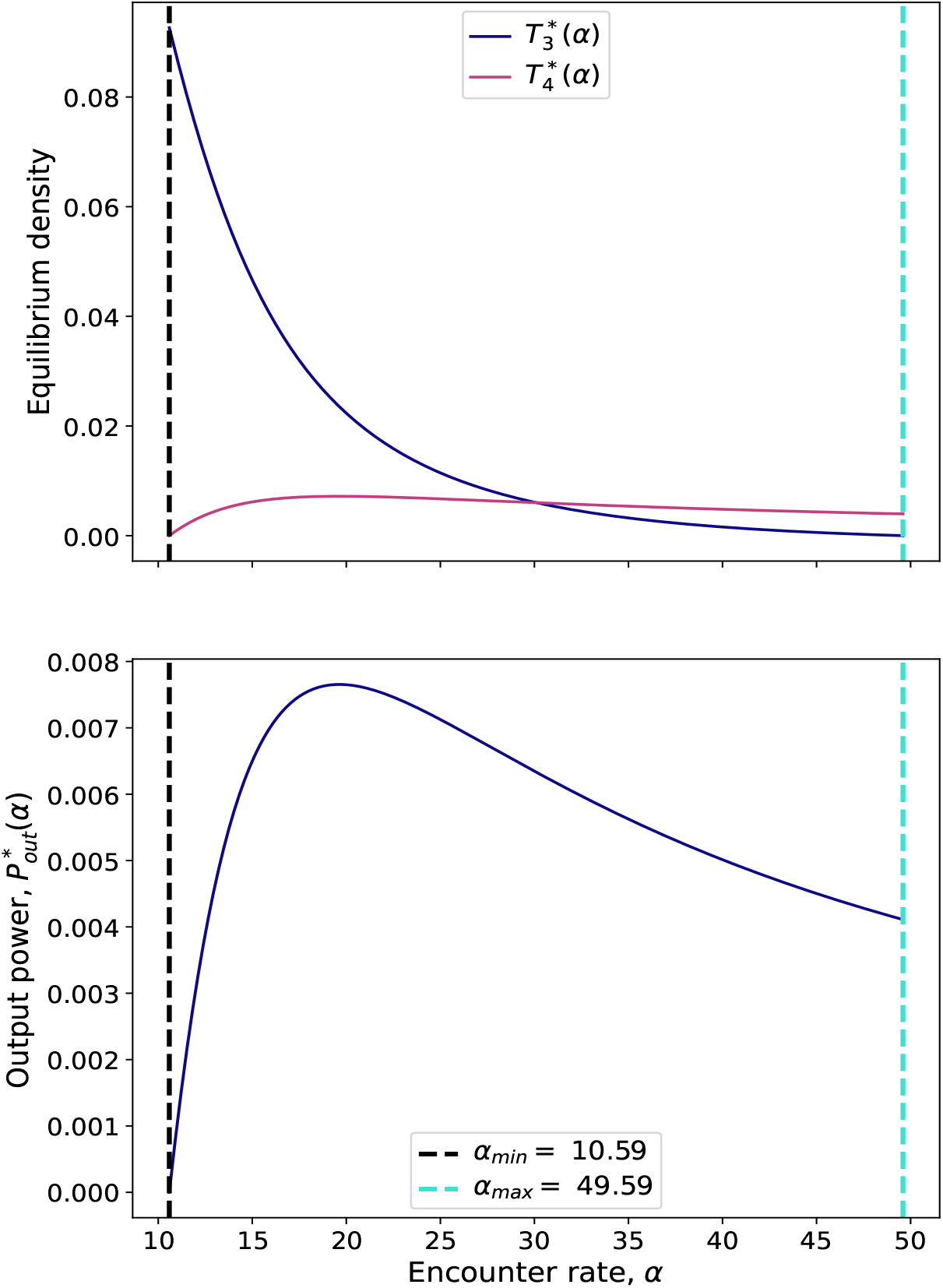
Four-trophic-level system with non-omnivory. Maximum output power no longer coincides with extinction of the intermediate species 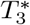. Parameter values are *a*_1_ = 1, *a*_2_ = 2, *a*_3_ = 4, *a*_4_ = 9, *k*_12_ = 1.414, *k*_23_ = 2, *k*_34_ = 3, *k*_24_ = 3, *ε* = 0.1, *r*_1_ = 5, *d*_2_ = 0.1, *d*_3_ = 0.1, and *d*_4_ = 1.

For the corresponding basal-omnivory system, the equations become

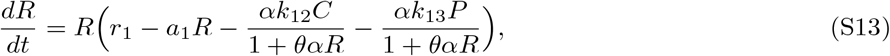

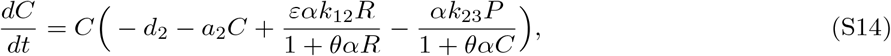

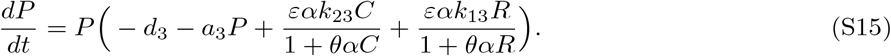

Because exact analytical expressions for feasible equilibria are not available in these cases, we compute all quantities numerically. To estimate the probability of feasibility, we sample large numbers of growth-rate vectors and compute the fraction that yield positive equilibria.

**Figure S10:**
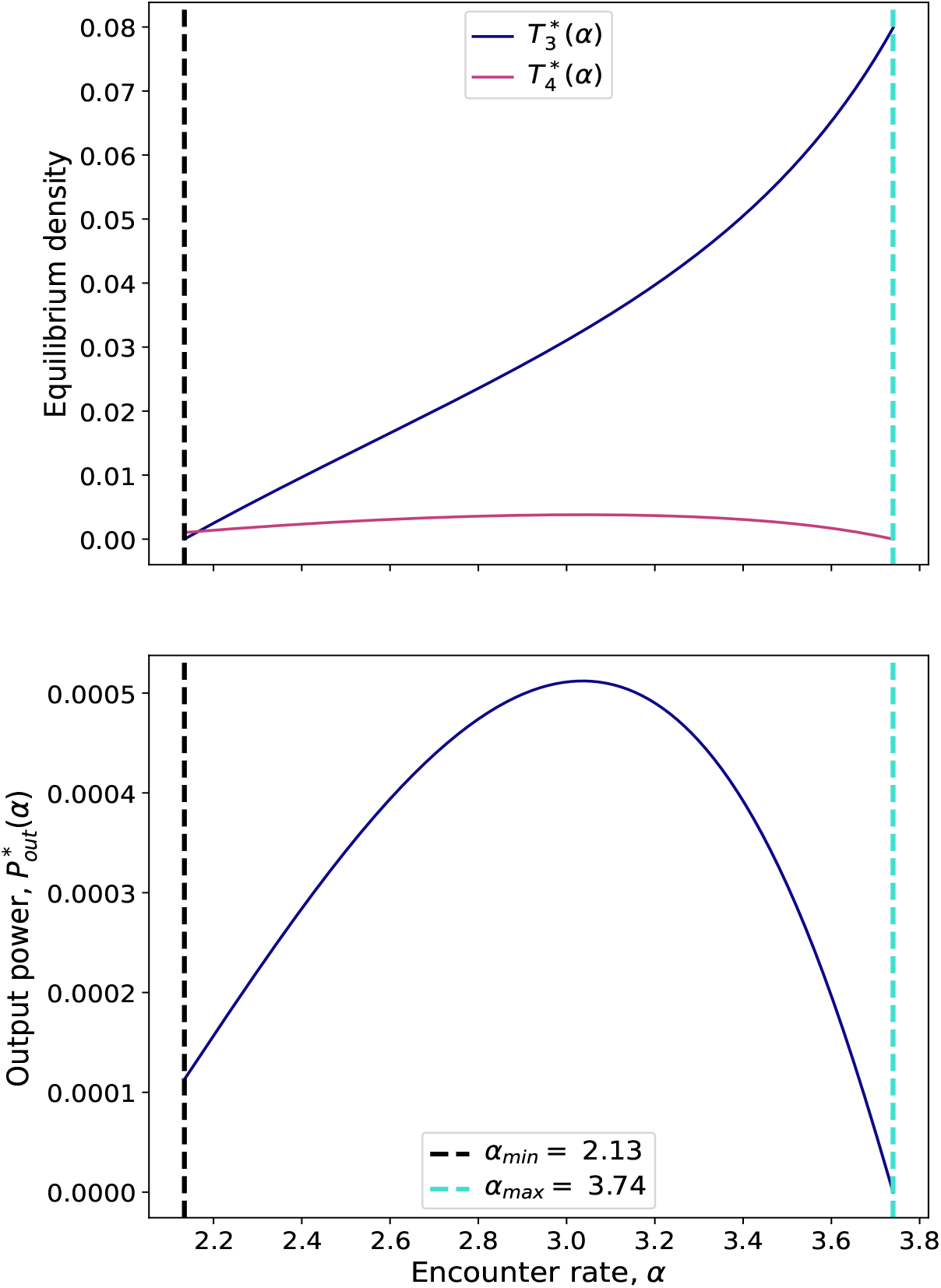
Four-trophic-level system with both basal and non-omnivory. The qualitative behaviour resembles the non-basal case, with maximum output power decoupled from extinction of the intermediate trophic species. Parameter values are *a*_1_ = 1, *a*_2_ = 2, *a*_3_ = 4, *a*_4_ = 9, *k*_12_ = 1.414, *k*_23_ = 2, *k*_34_ = 3, *k*_13_ = 2, *k*_24_ = 3, *ε* = 0.1, *r*_1_ = 2, *d*_2_ = 0.1, *d*_3_ = 0.7, and *d*_4_ = 0.1.

The numerical results preserve the qualitative distinction between trophic chains and omnivory. In the chain, the feasibility fraction and output power increase together. In the basal-omnivory case, by contrast, maximum output power again coincides with the extinction threshold of the herbivore-like intermediate species.

**Figure S11:**
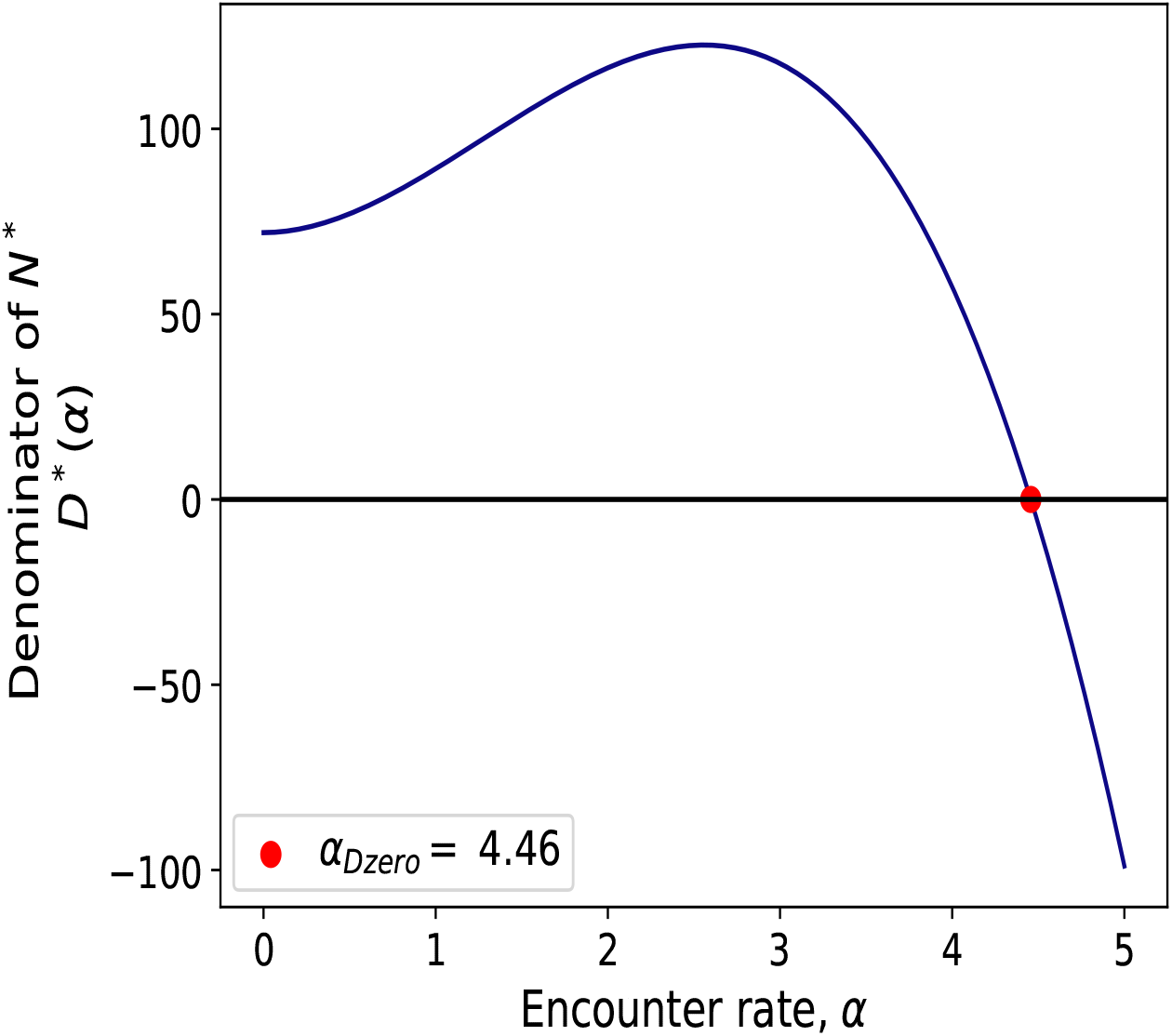
For the four-trophic-level system with both basal and non-omnivory, we plot the denominator *D*\*(*α*) of the equilibrium densities. *α*_*Dzero*_ also corresponds to the point of minimum Ω(*α*) as in Figure 2. Parameter values are *a*_1_ = 1, *a*_2_ = 2, *a*_3_ = 4, *a*_4_ = 9, *k*_12_ = 1.414, *k*_23_ = 2, *k*_34_ = 3, *k*_13_ = 2, *k*_24_ = 3, *ε* = 0.1, *r*_1_ = 2, *d*_2_ = 0.1, *d*_3_ = 0.7, and *d*_4_ = 0.1.

**Figure S12:**
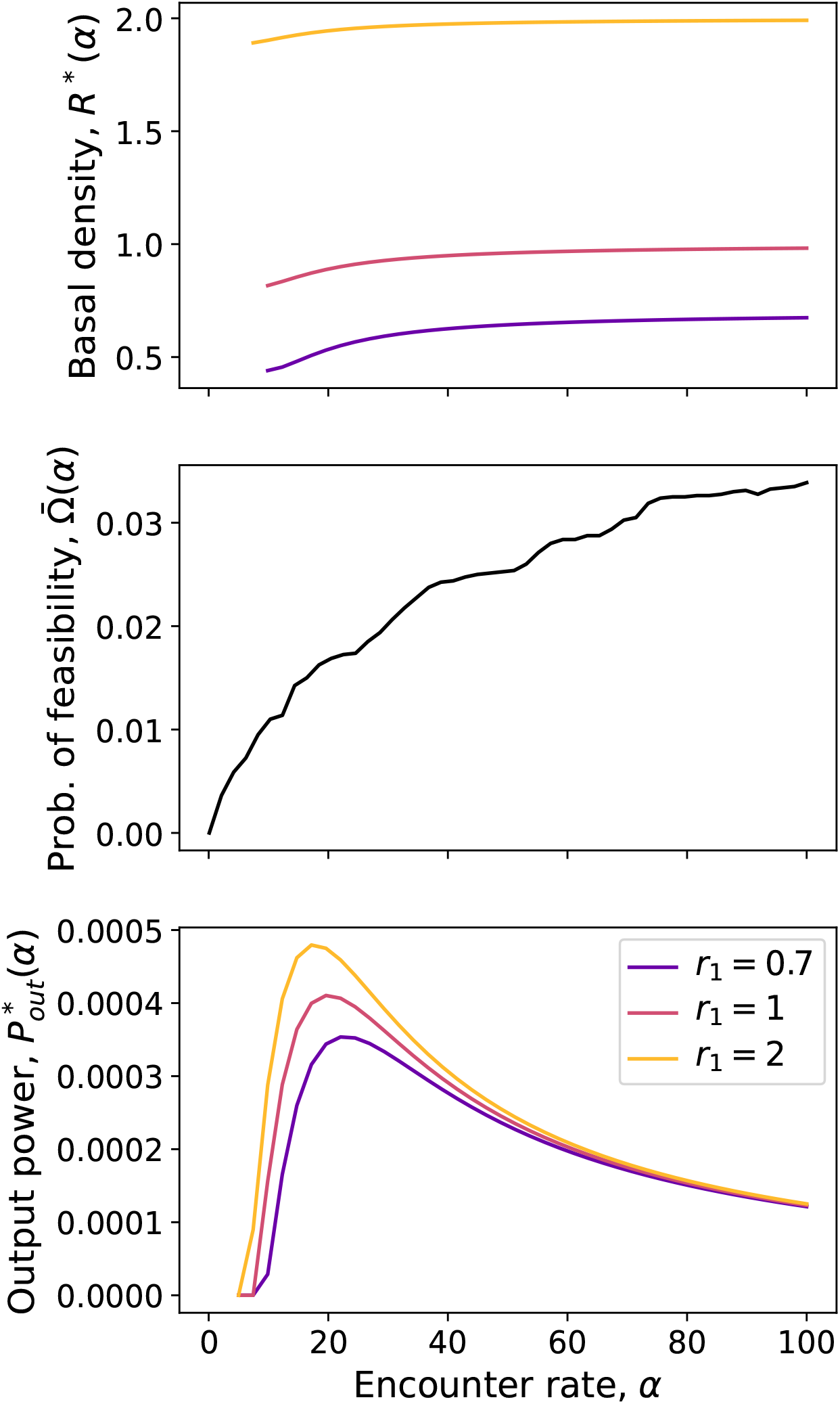
Three-species trophic chain with type II functional response. The figure shows the equilibrium density of the basal species *R*\*(*α*), the sampled feasibility fraction 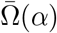, and output power 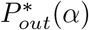 as functions of *α*. Parameter values are *a*_1_ = 1, *a*_2_ = 4, *a*_3_ = 9, *k*_12_ = 2, *k*_23_ = 3, *θ* = 0.5, *ε* = 0.1, *d*_2_ = 0.1, and *d*_3_ = 0.1.

**Figure S13:**
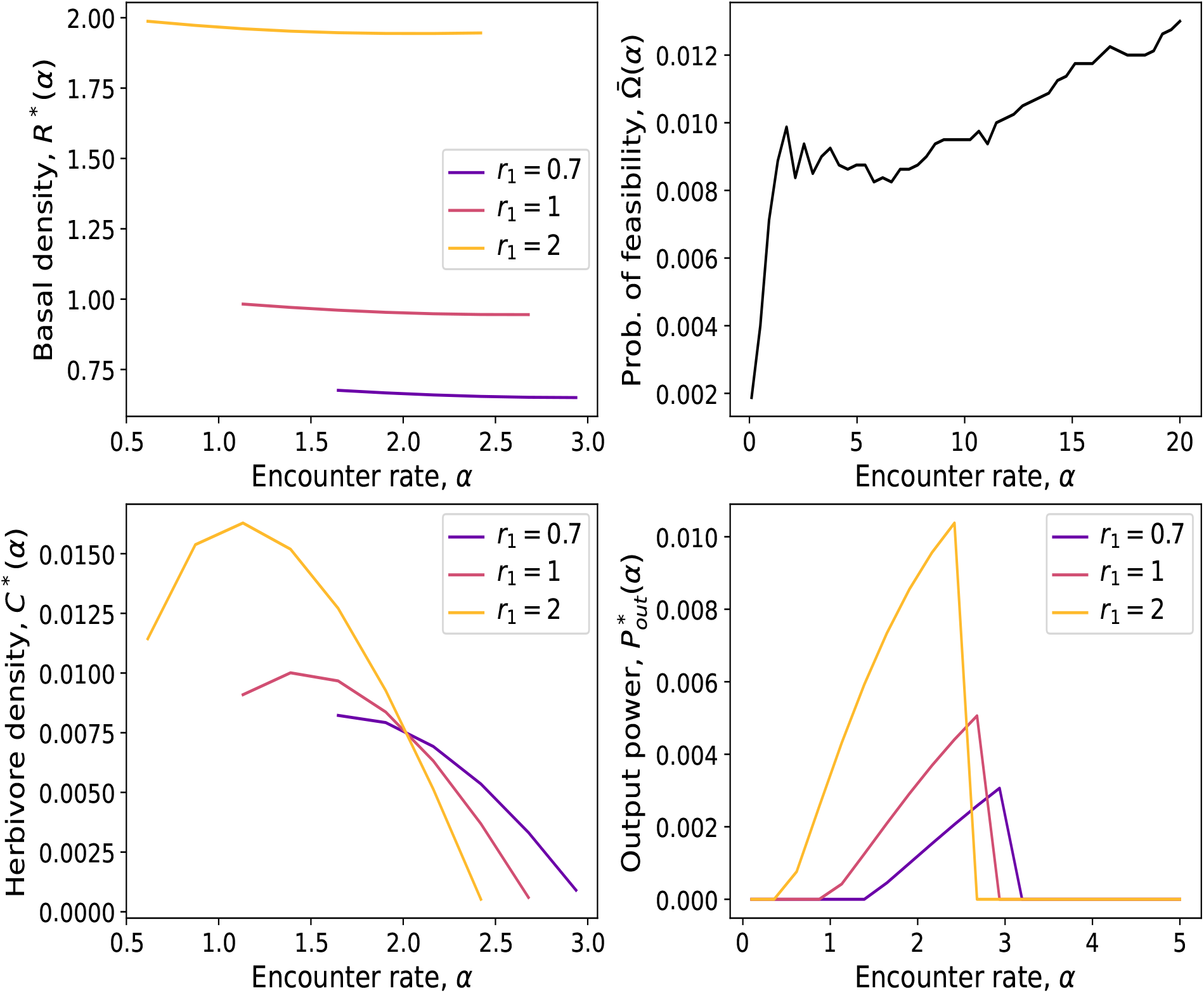
Three-species basal-omnivory system with type II functional response. The figure shows *R*\*(*α*), *C*\*(*α*), the sampled feasibility fraction 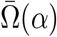, and output power 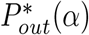 as functions of *α*. Parameter values are *a*_1_ = 1, *a*_2_ = 4, *a*_3_ = 9, *k*_12_ = 2, *k*_23_ = 3, *k*_13_ = 3, *θ* = 0.5, *ε* = 0.1, *d*_2_ = 0.1, and *d*_3_ = 0.2.

